# Ectoine production through a marine methanotroph-microalgae culture allows complete biogas valorization

**DOI:** 10.1101/2024.09.20.613850

**Authors:** Patricia Ruiz-Ruiz, Patricia Mohedano-Caballero, Jo De Vrieze

**Affiliations:** Center for Microbial Ecology and Technology (CMET), Ghent University, Frieda Saeysstraat 1, B-9052 Gent, Belgium; Centre for Advanced Process Technology for Urban Resource recovery (CAPTURE), Frieda Saeysstraat 1, B-9052 Gent, Belgium

**Keywords:** Biogas, ectoine, enrichment, GHG, methane, methanotrophs, microalgae

## Abstract

Methanotrophs have recently emerged as a promising platform for producing bio-based chemicals, like ectoine, from biogas, offering an economical alternative to glucose. However, most studies have focused solely on CH_4_ consumption, often overlooking the CO_2_, which is both produced by methanotrophs and present in biogas, despite its potential as a carbon source for microorganisms, such as microalgae. In this study, marine methanotrophic-microalgal cultures were enriched from environmental samples collected at the North Sea coast to explore ectoine production from both CH_4_ and CO_2_ in biogas. The sediment-derived culture exhibited the highest CH_4_ removal efficiency and CO_2_ uptake, and was selected for further experiments. The culture was primarily composed of *Methylobacter marinus*, *Methylophaga marina*, and the microalga *Picochlorum oklahomensis*. Gas consumption, growth, and ectoine production were evaluated under varying salinity levels and osmotic stress. The NaCl concentrations above 6% negatively impacted CH_4_ oxidation and inhibited ectoine synthesis, while osmotic shocks enhanced ectoine accumulation, with a maximum ectoine content of 51.3 mg_ectoine_ g_VSS_^−1^ at 4.5% NaCl. This study is the first to report ectoine production from methanotroph-microalgal cultures, showing its potential for biogas valorization into high-value bio-based chemicals, like ectoine, marking a significant step toward sustainable biogas utilization.

**Highlights:**

- Methanotroph-microalgae cultures can valorize CH_4_ and CO_2_ from biogas into ectoine
- NaCl concentrations above 6% reduced CH_4_ uptake and inhibited ectoine synthesis
- Osmotic shock at 4.5% NaCl enhanced ectoine accumulation
- First report of ectoine production using methanotroph-microalgal cultures

## 1. Introduction

As the world shifts towards a sustainable circular bioeconomy, the recovery of resources from waste streams into valuable goods becomes increasingly essential. Biogas production and utilization as a renewable energy source have gained attention over the past decade. Biogas, produced through anaerobic digestion (AD) of organic matter, consists mainly of 50−70% methane (CH_4_) and 30−50% carbon dioxide (CO_2_). The primary goal in anaerobic digesters is to generate CH_4_ for power or heat, with combined heat and power (CHP) units being the most common method of biogas valorization (Acosta and De Vrieze, 2019). However, small-scale decentralized biogas digesters frequently struggle with unstable feedstock and poor management, leading to low conversion efficiencies (20-40%) and low CH_4_ concentrations. While a minimum CH_4_ content of 35-40% is needed for CHP systems to be economically viable, upgrading biogas to meet this threshold can be costly, and CHP installation is often impractical for decentralized operations (Acosta and De Vrieze, 2019). As a result, biogas is typically burned with limited heat recovery instead of being upgraded for electricity. Additionally, electricity is a relatively low-value product, reducing the appeal of this approach, which can lead to biogas flaring or, in the worst case, venting, both contributing to greenhouse gas (GHG) emissions. Therefore, converting CH_4_ and CO_2_ into higher-value products is crucial to justify investments in biogas capture and utilization.

In recent years, methanotrophs have been recognized as a promising platform for producing valuable products from biogas (Cantera et al., 2016; Strong et al., 2016). One example is ectoine (1,4,5,6-tetrahydro-2-methyl-4-pyrimidine carboxylic acid), an osmolyte known for protecting cells against osmotic dehydration and stabilizing enzymes and nucleic acids in highly saline environments (Khmelenina et al., 1999). Ectoine has become a high-demand chemical, due to its wide applications in biotechnology, cosmetics, and medicine, representing a multibillion-dollar market with an annual demand of about 15,000 tons and a retail price of approximately 1,000 USD per kilogram (Liu et al., 2021).

Several halotolerant methanotrophs, including *Methylomicrobium alcaliphilum*, *Methylobacter marinus*, *Methylomicrobium kenyense*, *Methylophaga thalassica*, *Methylophaga alcalica*, and *Methylarcula marina*, have the ability to produce and accumulate ectoine (Cantera et al., 2016; Khmelenina et al., 1999). Traditionally, ectoine is industrially produced by the halotolerant heterotrophic proteobacterium *Halomonas elongata* through a process known as "bacterial milking". This method involves stimulating ectoine biosynthesis with a high-salt medium, followed by an osmotic shock with a low-salt medium to quickly release the synthesized ectoine (Liu et al., 2021). While this approach is effective, it is expensive, due to the need for high-quality carbon sources, such as glucose, sodium glutamate, and yeast extract. The idea of using biogas as a feedstock presents a potentially more economical and attractive alternative to these costly substrates, warranting further investigation. However, current ectoine yields from methanotrophic cultures are still lower to those achieved through industrial processes. Among methanotrophs, the genetically engineered *Methylomicrobium alcaliphilum* 20Z is the most efficient ectoine producer, with a yield of 0.111 g_ectoine_ g_biomass_^−1^ using CH_4_, whereas sugar-fermenting microbes can produce up to 0.22 g_ectoine_ g_biomass_^−1^ using glucose (Cho et al., 2022). Ongoing research is focused on optimizing various parameters to improve ectoine production by methanotrophs, including NaCl content, pH, temperature, essential micronutrients, bioreactor operation strategies, and genetic modifications (Carmona-Martínez et al., 2021; Cho et al., 2022; Rodero et al., 2022).

Current studies have primarily focused on using methanotrophs alone to consume CH_4_ in biogas for ectoine production, often neglecting the other important component, CO_2_, present in both biogas and produced via methanotrophic activity. This oversight misses the opportunity to utilize CO_2_ as a carbon source for other microorganisms, such as microalgae, resulting in the underutilization of valuable carbon. To address this gap, the combined use of methanotrophs and microalgae, a strategy referred to as "methalgae" by Van Der Ha et al. (2011), offers a solution for maximizing the valorization of both CH_4_ and CO_2_ in biogas. This approach integrates CH_4_ oxidation with oxygenic photosynthesis, enabling the efficient use of both gasses within a single system. The synergy between methanotrophs and microalgae has yielded several successful outcomes, including nutrient recovery from wastewater, reduction of GHG emissions, and the production of valuable compounds, such as single-cell protein, biopolymers, and lipids (Li et al., 2022; Rasouli et al., 2018; Ruiz-Ruiz et al., 2024, 2020; Van Der Ha et al., 2012).

Although data on ectoine accumulation in microalgae remain uncertain, with some reports indicating minimal levels of this osmolyte in certain microalgal species (McParland et al., 2021), the stimulation of ectoine production in co-cultures with ectoine-producing bacteria has been observed (Di Costanzo et al., 2021; Fenizia et al., 2021, 2020), though not in methanotrophs thus far. Additionally, several studies have highlighted the benefits of combining microalgae with methanotrophic cultures from a bioprocess perspective (Hill et al., 2017; Li et al., 2022; Ruiz-Ruiz et al., 2024; Yun et al., 2024). These benefits include enhanced process efficiency through the provision of oxygen and essential growth factors, mitigation of toxicity from byproducts, such as organic acids, and regulation of culture pH through the coordinated activity of each microbial group. This combination also has the potential to achieve a zero-GHG process, while producing valuable compounds (Ruiz-Ruiz et al., 2024).

In this work, we propose a novel value chain for biogas by utilizing both CH_4_ and CO_2_ within a single system to produce the high-value metabolite ectoine. The primary objective was to enrich ectoine-producing mixed methanotrophic-microalgal cultures derived from saline environments, using synthetic biogas as the sole carbon source. The effects of varying salinity levels and osmotic shocks on gas consumption, growth, and ectoine content of the methalgae culture were evaluated. This approach marks a substantial advancement in harnessing the potential of methanotrophic-microalgal cultures to effectively valorize biogas for ectoine production, a potential that has not been explored to date.

## 2. Methods

### 2.1 Marine methalgae enrichments and culture conditions

Three environmental sampling points were sourced for methanotrophic bacteria and microalgae from the Zwin Natuur Park (**Fig. S1**), a nature reserve on the North Sea coast at the Belgian-Dutch border (51.3580N, 3.3459E). The samples, named Sediment, North Sea, River, and a Combination of these three, were enriched in 120 mL transparent glass bottles sealed with butyl septa and aluminum caps.

Initially, 2 grams of combined soil and water from the sampling points were added to 20 mL of sterile mineral salt medium (MSM) containing (in g L^−1^): KNO_3_, 1.0; MgSO_4_·7H_2_O, 1.0; CaCl_2_·2H_2_O, 0.15; FeNaEDTA, 0.005; Na_2_HPO_4_·12H_2_O, 0.717; KH_2_PO_4_, 0.272; and 1 mL of the following trace elements (in g L^−1^): Na_2_EDTA·2H_2_O, 0.5; FeSO_2_·7H_2_O, 0.2; H_3_BO_3_, 0.03; CoCl_2_·6H_2_O, 0.02; ZnSO_4_·7H_2_O, 0.01; MnCl_2_·4H_2_O, 0.003; NaMoO_4_·2H_2_O, 0.003; NiCl_2_·6H_2_O, 0.002; CuSO_4_·5H_2_O, 2.5, pH 6.8. A NaCl concentration of 3% was added to the culture medium for all enrichments. Pure CH_4_ (>99%) was filtered through 0.2 μm pore size filters and injected into the headspace using a 60 mL syringe to achieve a concentration of ∼10% CH_4_ in air (average mass of 5.5 mg CH_4_). To prevent overpressure in the glass bottles, an equal volume of air was removed immediately before the CH_4_ injection. No CO_2_ was added to the enrichments to ensure that the CO_2_ produced via methanotrophy was the sole carbon source for the photosynthetic microorganisms. The headspace gas composition was renewed every 2 days, with initial and final CH_4_, O_2_, and CO_2_ concentrations monitored by gas chromatography (GC-TCD). Every two weeks, 2 mL aliquots from the previous enrichment were sequentially transferred to new sterile glass bottles containing 18 mL of fresh MSM with 3% NaCl, over a total period of 10 weeks. The bottles were incubated in an orbital shaker at 125 rpm in a temperature-controlled room (28 ± 2 °C). Full-spectrum LED lights were placed above the orbital shaker to provide an average irradiance of 120 μmol m^−2^ s^−1^. Enrichments were performed in duplicate. The methalgae enrichment that presented the highest CH_4_ removal efficiencies, as calculated according to section 2.6, was selected for subsequent experiments.

### 2.2 Growth of methalgae enrichment at different NaCl concentrations

The selected enriched methalgae culture was pre-grown in a 1.2 L Schott bottle with a plastic screw cap and butyl septa for 7 days to produce sufficient inoculum for this experiment. The culture conditions adhered to those described in Section 2.1, using MSM with 3% NaCl. To study the effects of varying NaCl concentrations on gas consumption, growth, and ectoine accumulation, the biomass was centrifuged (7,745 xg) and washed three times with sterile PBS before being resuspended in 200 mL of sterile MSM containing 0%, 3%, 6%, and 9% NaCl concentrations. The experiments were conducted in 0.6 L transparent gas-tight bottles, set up in biological duplicates with 0.2 L of liquid culture. The initial biomass concentration was 0.4 ± 0.01 g_VSS_ L^−1^. The bottles were incubated on an orbital shaker at 125 rpm in a temperature-controlled room maintained at 28 ± 2 °C. Full-spectrum LED lights were positioned above the orbital shaker to provide an average irradiance of 120 μmol m^−2^ s^−1^. The headspace of each bottle contained diluted CH_4_ in air (1:1.5 CH_4_:O_2_ volume ratio) giving an average mass of 18 mg CH_4_, which was renewed every 48 h. Initial and final gas samples were collected to measure CH_4_, O_2_, and CO_2_ concentrations by GC-TCD, as detailed in section 2.6. Liquid samples were taken to assess total suspended solids (TSS) and volatile suspended solids (VSS), cell counts via flow cytometry (FCM), and ectoine content. The experiment was conducted over a period of 14 days.

### 2.3 Osmotic shock test

To investigate the effect of varying osmotic shocks on gas uptake, growth, and ectoine accumulation in the methanotrophic-microalgal culture, an experimental setup was designed where salinity levels were adjusted over time. The methalgae culture was pre-grown in a 1.2 L Schott bottle with a plastic screw cap and butyl septa for 7 days, following the culture conditions described in Section 2.1. This pre-growth phase, using MSM with 3% NaCl, ensured sufficient inoculum for the experiment. The main experiments were conducted in 0.6 L transparent gas-tight bottles, with biological duplicates consisting of 0.15 L of liquid culture and 0.45 L of headspace. The initial biomass concentration was 0.45 ± 0.06 g_VSS_ L^−1^. Osmotic shocks were administered at different time points, as detailed in **Table 1**. Before each osmotic shock, the biomass was sedimented by centrifuging the entire culture in 50 mL sterile tubes at 7,745 xg for 10 min, repeating the process until all of the culture had been fully centrifuged. The supernatant was then discarded, and the pellet was resuspended in 150 mL of fresh sterile MSM. This medium replacement process was conducted in a laminar flow cabinet and took approximately 2.5 h to complete before all cultures were fully restored. The headspace of each reactor contained a gas mixture of diluted synthetic biogas (1:1.5:0.6 CH_4_:O_2_:CO_2_ volume ratio), giving an average mass of 36 mg CH_4_, which was renewed every 24 h. Initial and final gas samples were taken to monitor CH_4_, O_2_, and CO_2_ concentrations by GC-TCD. Liquid samples were also collected to measure TSS/VSS, cell counts via FCM, and ectoine content. The experiment was carried out over 5 days.

**Table 1.** Percentage of NaCl over the experimental time in the three osmotic shock tests.

| Test |  | Time (hours) |  |  |  |
| --- | --- | --- | --- | --- | --- |
|  |  | 0 | 48 | 72 | 96 |
| OS1 | NaCl (%) | 3% | 4.5% | 6% | 3% |
| OS2 | NaCl (%) | 3% | 4.5% |  | 3% |
| OS3 | NaCl (%) | 3% | 6% |  | 3% |

### 2.4 Microbial composition analysis

The DNA extraction from the enriched culture was performed using the PowerSoil DNA Isolation Kit, following the manufacturer’s instructions. The extracted genomic DNA was then sent to LGC Genomics GmbH (Berlin, Germany) for library preparation and sequencing on the Illumina MiSeq platform using V3 chemistry. The V4 region of the 16S rRNA gene was targeted with primers 341F and 785Rmod (Klindworth et al., 2013) to identify bacterial domains, while the V4 region of the 18S rRNA gene was targeted with primers TAReuk454FWD1 and TAReukREV3 (Stoeck et al., 2010) to identify eukaryotic microalgae. The amplicon sequence data was processed using the DADA2 R package (Callahan et al., 2016). Primer sequences were removed, and reads were truncated at a quality score cut-off (truncQ=2). Additional filtering eliminated reads with ambiguous base calls or high expected errors (maxEE=2.2). After dereplication, unique reads were denoised using the DADA error estimation and selfConsist sample inference algorithms. The error rates were inspected and, once approved, the denoised reads were merged. Amplicon sequence variants (ASVs) were obtained after chimera removal and used for taxonomy assignment with the Naive Bayesian Classifier and the DADA2-formatted Silva v138.1 database. The amplicon sequence data have been deposited in the NCBI Sequence Read Archive (SRA) database under BioProject ID PRJNA1161840, with accession numbers SRX26108882 and SRX26108883.

### 2.6 Analytical procedures

#### Gas chromatography

The gas phase composition was analyzed with a Compact GC4.0 (Global Analyzer Solutions, Breda, The Netherlands), equipped with a Molsieve 5A pre-column and Porabond Q column (CH_4_, O_2_, H_2_ and N_2_) and a Rt-Q-bond pre-column and column (CO_2_, N_2_O and H_2_S). Gas concentrations were determined using a thermal conductivity detector (TCD) with a minimum detection limit of 100 ppm_v_ for each gas.

#### Biomass (TSS/VSS) and pH

Total (TSS) and Volatile (VSS) Suspended Solids were measured according to Standard Methods (APHA, 2012) using glass fiber membrane filters of 0.3 μm pore size. Biomass estimation was reported as VSS instead of TSS, due to the observation that high NaCl concentrations affected the accuracy of TSS measurements. The pH measurement of the liquid samples was carried out manually with a micro pH electrode (Consort C3210 Multi-parameter analyzer, Belgium).

#### Ectoine extraction and measurement

For intracellular ectoine measurement, biomass was concentrated threefold by centrifugation of 2 mL of the culture broth at 21,130 xg for 5 min. The resulting pellets were stored at −20 °C until further ectoine extraction. An aliquot of 1.5 mL of a solution of 80% ethanol and 20 ± 5 mg of 0.1 mm diameter zirconia/glass beads (Biospec Products, Belgium) was added to the tube, followed by cell disruption in a PowerLyzer (Mo Bio Laboratory Inc., USA) at 2000 rpm for 10 min. The supernatant was kept at 4 °C overnight. The suspension was then centrifuged at 21,130 xg for 5 min and filtered through 0.22 µm filters before analysis by HPLC-UV. For extracellular ectoine content determination, the supernatant obtained after sample centrifugation was filtered through a 0.22 μm filter prior to measurement. Commercial ectoine (≥95.0%, Sigma Aldrich) was used to prepare the calibration curve. Ectoine was analyzed by HPLC-UV using an Aminex HPX-87C column (Bio-Rad Laboratories, USA), with CaCl_2_ (5 mM) as the eluent and detection at 210 nm (Onraedt et al., 2004). The flow rate was 0.5 mL min^−1^, the column temperature was 85 °C, and the runtime was 52 min. The specific ectoine yield (in mg_ectoine_ g_vss_^−1^) was calculated using the biomass concentration estimated through the VSS content.

#### Flow cytometry

To measure the total number of cells, a flow cytometer (Attune NxT FCM) was used. A 180 μL sample, previously sonicated for 3 min to disaggregate culture granules, was placed in 96-well plates in triplicate. Dilutions were made to ensure that the samples contained no more than 10^6^ cells mL^−1^. Then, 1% volume of Sybr Green stain was added to the sample and incubated at 37 °C for 30 min (Van Nevel et al., 2013). Control samples of the filtered medium (0.22 μm) at every NaCl concentration were used for background removal and cell counting. Data analysis was performed using FCS Express.

### 2.7 Calculations and statistical analysis

Removal efficiency (RE, %) of gases was calculated using Equation (1), where *C_gas initial_* is the initial gas concentration and *C_gas final_* is the final gas concentration between gas replacements. For gas calculations atmospheric pressure of 1 atm and a temperature of 28 °C was considered.

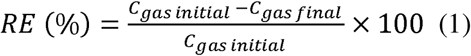

The statistical data analysis was performed using SPSS 20.0 (IBM, USA). The results are presented as the average ± standard deviation. For normally distributed and homoscedastic data, t-tests were used to compare the effects of different variables on experimental groups. The non-parametric Mann-Whitney U test was used for two-sample comparisons, and the Kruskal-Wallis test was used for comparisons involving more than two independent samples in case normality and/or homoscedasticity could not be confirmed. Differences were considered significant at p < 0.05.

## 3. Results

### 3.1 Enrichments performance

The headspace gas evolution for all enrichments is depicted in **Fig. 1**. The Sediment enrichment demonstrated higher and more consistent CH_4_ oxidation and CO_2_ production compared to the other three enrichments, achieving an average CH_4_ removal efficiency (RE) of 88 ± 16% throughout the incubation period (**Fig. 2a**). The calculated CO_2_ produced by methanotrophs from CH_4_ consumption in the Sediment enrichment (**Fig. S2a**) indicates that some of this CO_2_ was fixed by microalgae, as no CO_2_ was added to the headspace and none was retained in the medium (pH 6.5-6.8 throughout). Photosynthetic CO_2_ fixation produced O_2_, preventing the bottles from becoming anoxic (**Fig. 1**).

**Fig. 1.**
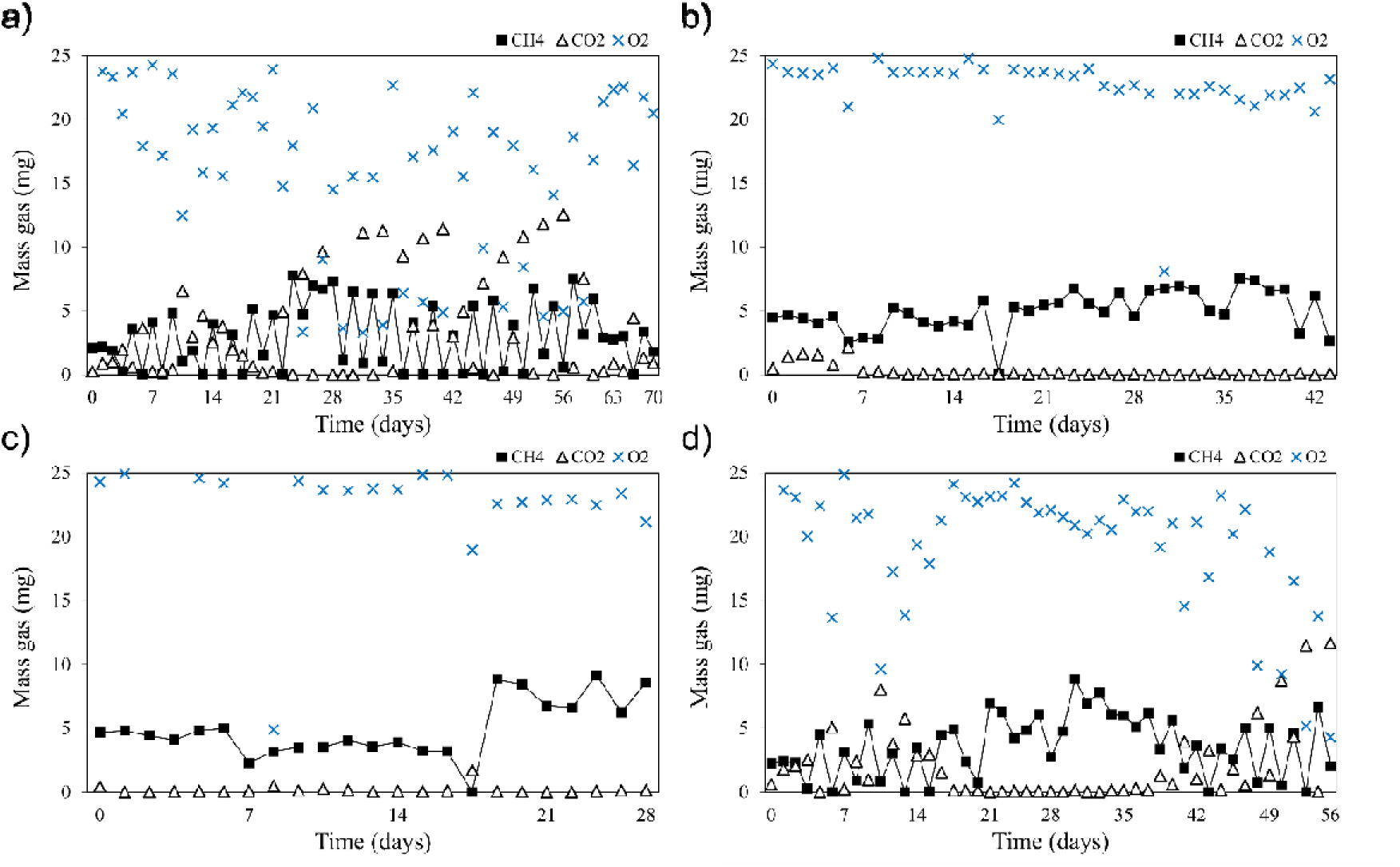
Headspace gas evolution of CH_4_, CO_2_, and O_2_ of the enrichments during the incubation period. a) Sediment. b) River. c) North Sea. d) Combination.

**Fig. 2.**
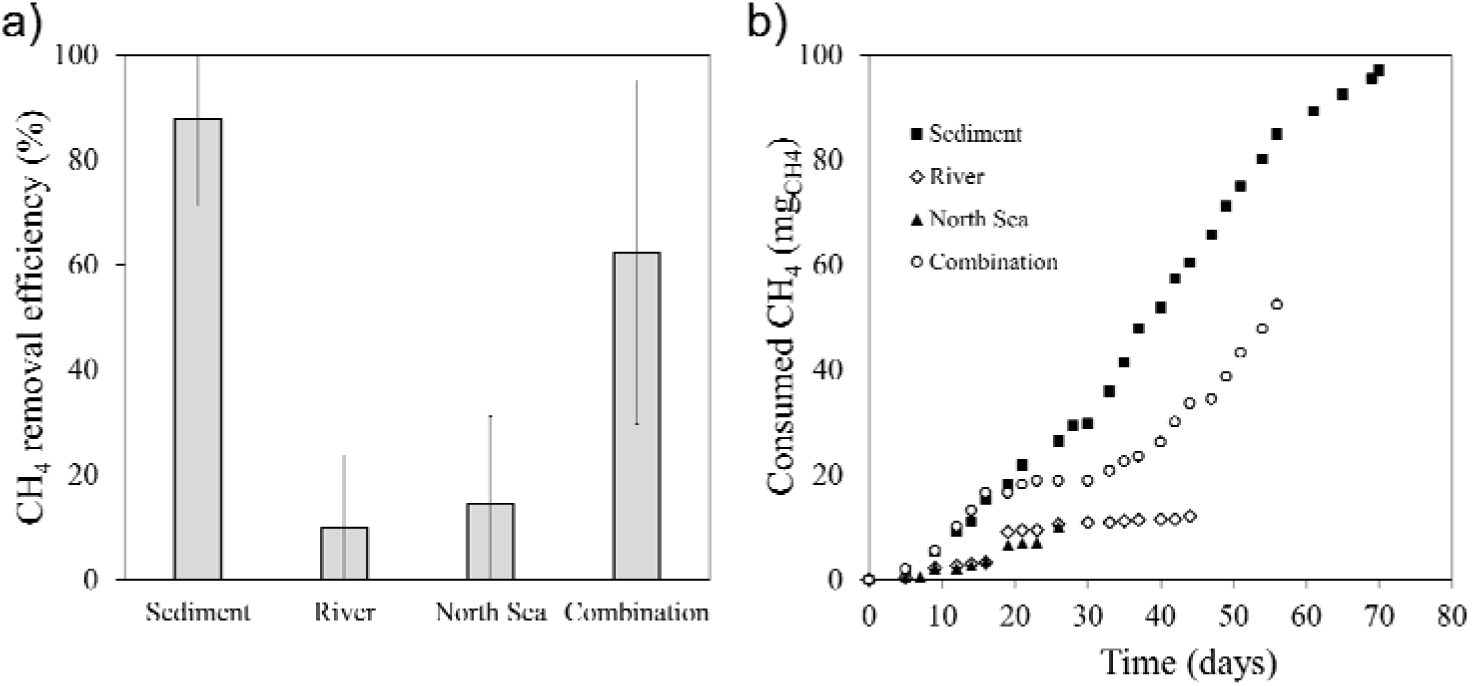
a) Average CH_4_ removal efficiency (%) for the four different enrichments during the entire incubation period. b) Cumulative CH_4_ consumption over time (mg_CH4_ consumed).

In contrast, the River and North Sea enrichments showed minimal CH_4_ oxidation, resulting in no O_2_ consumption (**Fig. 1**) and poor CO_2_ production (**Fig. S2**), with CH_4_ REs below 15% in both cases (**Fig. 2a**). The Combination enrichment displayed fluctuating gas production and consumption, with an average CH_4_ RE of 62 ± 33% (**Fig. 2a**). Significantly higher CH_4_ REs were observed in the Sediment and Combination enrichments compared to the North Sea and River enrichments (p < 0.0001) (**Fig. 2a**). By day 55, the Sediment enrichment had consumed 1.5 times more CH_4_ than the Combination enrichment, ultimately consuming nearly 100 mg of CH_4_ by the end of the experiment (**Fig. 2b**).

Enrichments with poor CH_4_ consumption were discarded earlier: North Sea after 28 days, River after 42 days, and Combination after 56 days. The Sediment enrichment was selected for further experiments due to its high CH_4_ uptake, CO_2_ production and consumption, and visible microalgal growth (**Fig. S3**).

### 3.2 Effect of NaCl concentration on gases consumption, growth, and ectoine production

After the 14-day experiment with different NaCl concentrations, tests with 0% and 3% NaCl showed significantly higher CH_4_ removal efficiencies (REs) compared to the 6% and 9% NaCl tests, which showed minimal CH_4_ consumption (p < 0.0001) (**Fig. 3a**). The 0% NaCl test achieved the highest CH_4_ RE (100 ± 0%), followed by 3% NaCl (92 ± 17%), with 100 ± 5 mg and 80 ± 3 mg of CH_4_ consumed, respectively. Higher CO_2_ consumption was also observed in these tests, attributed to increased microalgal activity (**Fig. S4**).

**Fig. 3.**
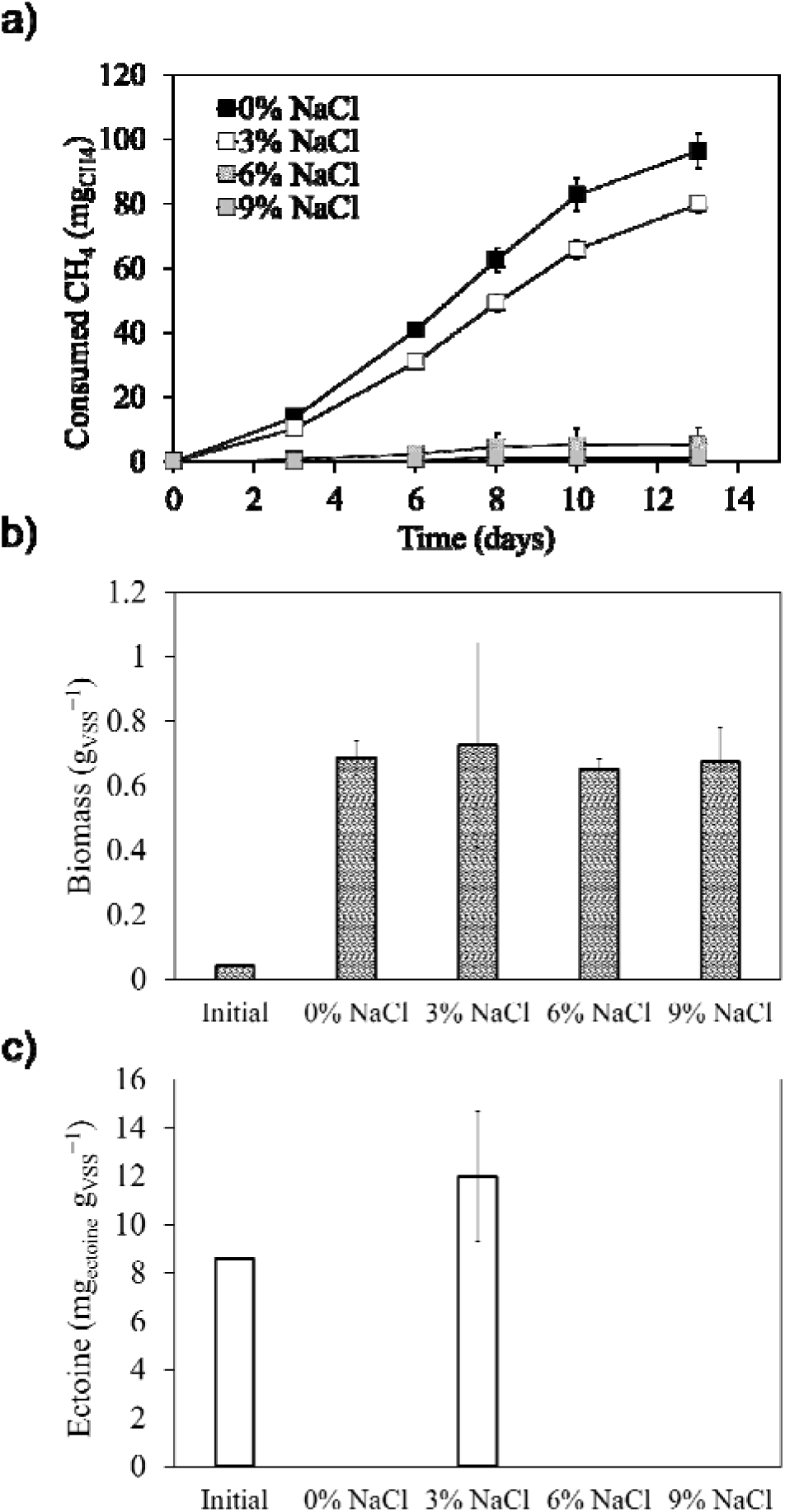
a) Cumulative CH_4_ consumption over time (mg_CH4_ consumed) for all NaCl concentrations. b) Average biomass concentration (g_VSS_ L^−1^) of the initial and final biomass samples. c) Average ectoine concentration (mg_ectoine_ g_VSS_^−1^). Error bars indicate the standard deviation of biological duplicates.

Photoinhibition of microalgae occurred on day 4 in one biological duplicate of the 3% and 9% NaCl tests due to excessive light exposure from a nearby experiment (**Fig. S5a**). This led to reduced CO_2_ consumption and O_2_ production (**Fig. S6**), though CH_4_ oxidation capacity remained unaffected in both the 3% (p = 0.2105) and 9% NaCl (p = 0.1793) tests.

A 17.5-fold biomass increase was observed across all experiments after 14 days, with an average biomass concentration of 0.68 ± 0.03 g_VSS_ L^−1^ (**Fig. 3b**). Microbial composition varied significantly, with microalgae making up 11% or less in the photoinhibited experiments, compared to 40-70% in the others (**Fig. S5b**). Bacteria dominated the remaining culture, and the photoinhibited tests helped refine the gating process for FCM analysis.

The initial inoculum had an intracellular ectoine concentration of 8.6 ± 0.1 mg_ectoine_ g_VSS_^−1^, which increased to 12.0 ± 2.7 mg_ectoine_ g_VSS_^−1^ in the 3% NaCl test after 14 days. No ectoine was detected in the biomass from the 0%, 6%, and 9% NaCl tests, nor in the supernatant at any NaCl concentration, indicating ectoine was not released into the medium (**Fig. 3c**).

### 3.3 Osmotic shock effect at different NaCl concentrations on gas consumption, growth, and ectoine accumulation

The effect of varying NaCl concentrations as osmotic shocks on CH_4_ RE was evaluated (**Fig. 4a**). Initially, all samples were grown in 3% NaCl for 7 days, achieving an average CH_4_ RE of 52 ± 3% before the osmotic shocks were applied. The first shock, increasing NaCl from 3% to 4.5% in OS1 and OS2, had no significant effect on CH_4_ RE (p = 0.33), which remained similar to the 3% NaCl medium. However, raising salinity to 6% in OS3 caused a substantial decrease in CH_4_ RE, about 6 times lower over the 24-48 h, consistent with previous findings.

**Fig. 4.**
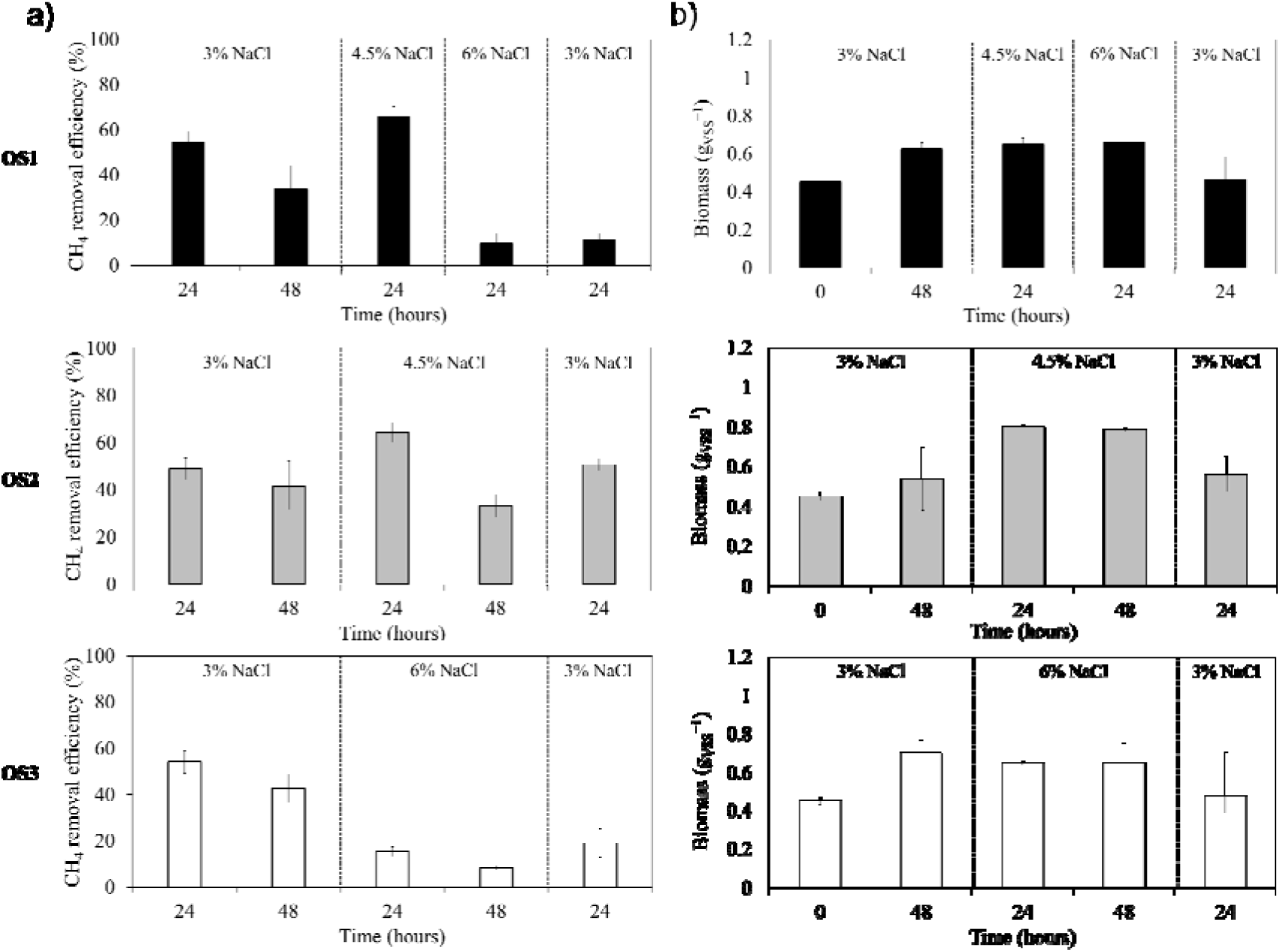
a) Average CH_4_ removal efficiency (%) per 24-h gas replacement for OS1, OS2, and OS3. b) Average biomass concentration (g_VSS_ L^−1^) for the initial and final biomass samples after each osmotic shock for OS1, OS2, and OS3. Error bars indicate the standard deviation of biological duplicates.

The second shock at 4.5% NaCl in OS2 caused a slight decrease in CH_4_ RE, though less severe than the drop seen at 6% NaCl in OS1. Despite minor improvements in CH_4_ RE after final incubation at 3% NaCl in OS1 and OS3, they did not reach the initial efficiency levels. In contrast, OS2’s RE after the final incubation at 3% NaCl returned to initial levels.

Higher NaCl concentrations (4.5% and 6%) also inhibited CO_2_ production compared to 3% NaCl (**Fig. S7**). At 6% NaCl, CH_4_ oxidation decreased, reducing O_2_ consumption and CO_2_ production by methanotrophs (**Fig. S8**). While OS1 and OS2 maintained stable activity at 4.5% NaCl, a significant reduction occurred at 6%, particularly in OS1, where O_2_, CO_2_, and CH_4_ levels remained constant, similar to OS3 during both shocks.

Biomass concentration increased after 48 h at 3% NaCl across all tests (**Fig. 4b**). After 24 h at 4.5% NaCl in OS1 and OS2, biomass continued to rise but was less pronounced at 6% NaCl. After re culturing at 3% NaCl, biomass content decreased in all tests.

Ectoine concentrations averaged 31.1 ± 1.5 mg_ectoine_ g_VSS_^−1^ after 48 h in 3% NaCl (**Fig. 5**), higher than in the salinity test, likely due to the increased abundance of methanotrophic and heterotrophic bacteria (**Table S1**). After the first osmotic shock at 4.5% NaCl, ectoine reached a maximum of 51.3 ± 1.1 mg_ectoine_ g_VSS_^−1^ in OS1. Ectoine levels remained stable during the first 24-48 h if salinity was unchanged, as seen in OS2 and OS3. However, ectoine levels decreased when salinity exceeded 4.5%, dropping from 51.3 ± 1.1 to 42.8 ± 3. mg_ectoine_ g_VSS_^−1^ in OS1. OS3 also showed reduced ectoine levels, three times lower than OS1 and OS2. Once cultures returned to 3% NaCl, ectoine levels slightly increased in all cases.

**Fig. 5.**
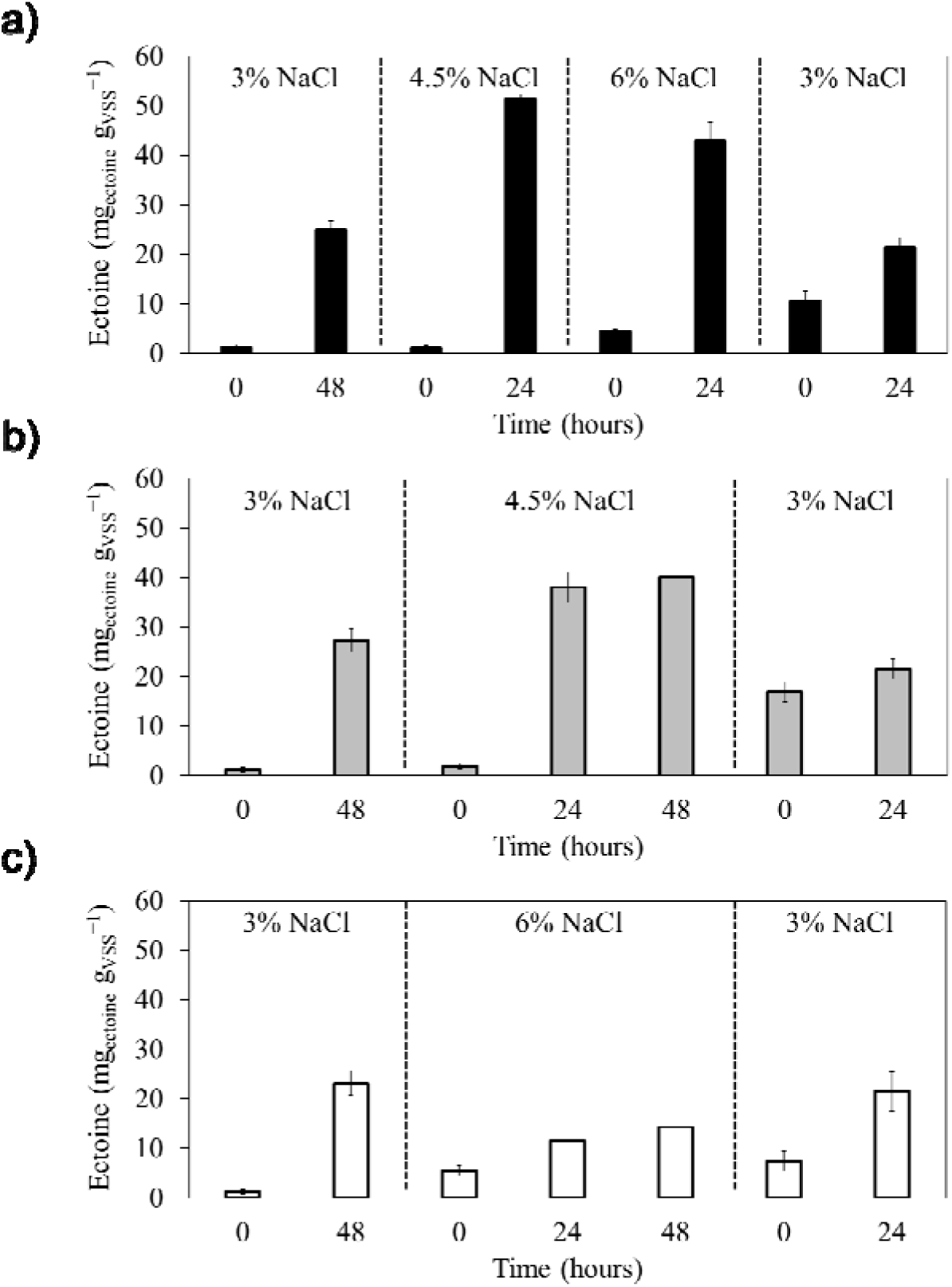
Intracellular ectoine concentration measured during the osmotic shock test (mg_ectoine_ g_VSS_^−1^). a) OS1. b) OS2 c) OS3. Error bars indicate the standard deviation of biological duplicates.

### 3.4 Structure of the enriched methalgae consortia

The 16S and 18S rRNA gene Illumina amplicon sequencing revealed the microbial composition and relative abundance of the Sediment enrichment (**Fig. 6**). *Picochlorum oklahomensis* (taxonomically revised to *oklahomense*) was the predominant photosynthetic eukaryotic microorganism, with a 97% relative abundance, followed by *Nannochloris* sp. with less than 1% relative abundance. The prokaryotic composition was more diverse, including various halophilic methylotrophic and heterotrophic microorganisms. The methanotroph *Methylobacter marinus/whittenburyi* and the methylotroph *Methylophaga marina* comprised 10% and 5% of the culture, respectively, followed by halophilic heterotrophic bacteria such as *Labrenzia* sp., *Labrenzia aggregata*, and *Hyphomonas adhaerens*.

**Fig. 6.**
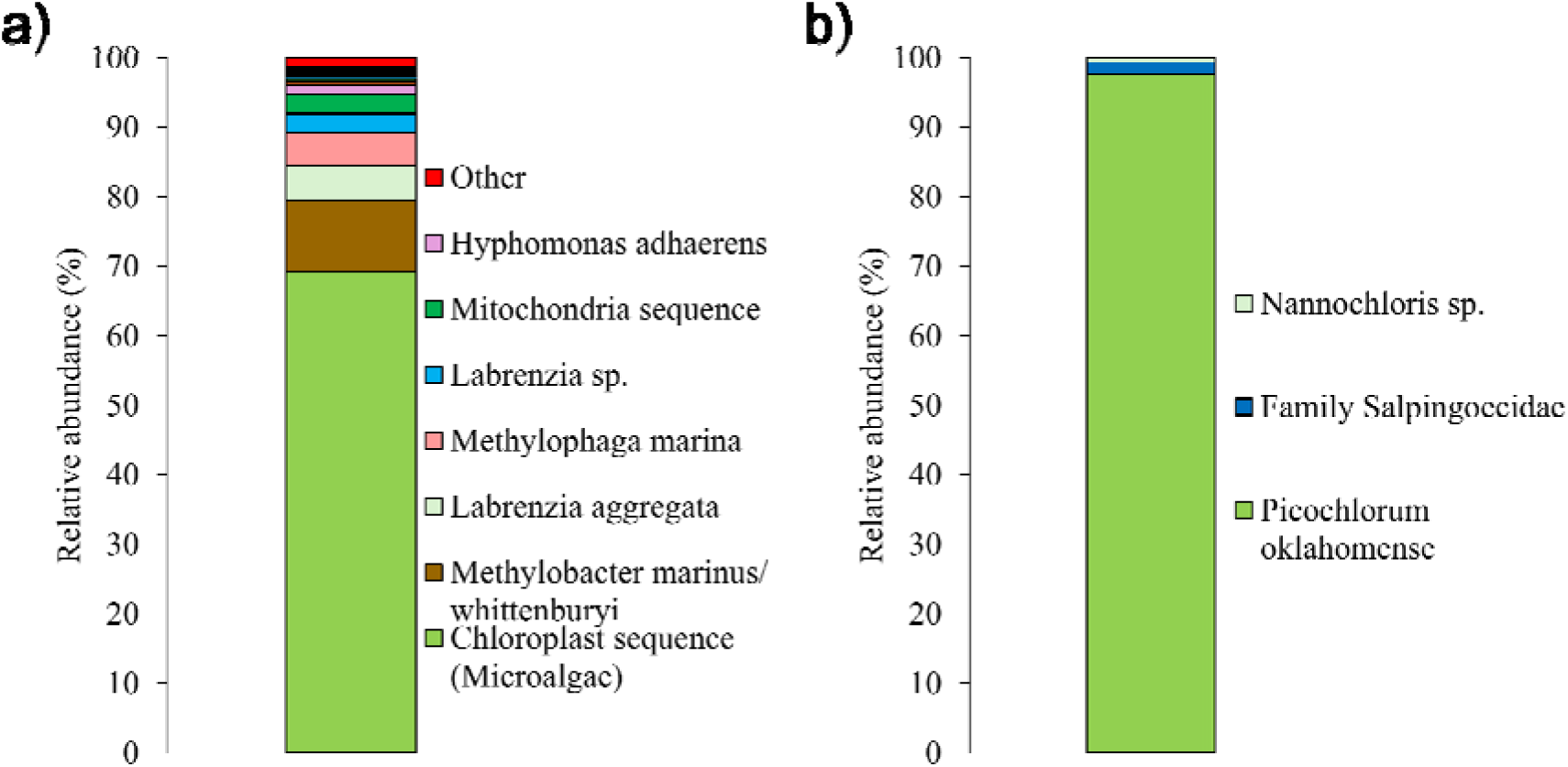
Relative abundance (%) of the prokaryotic and eukaryotic amplicon sequencing variants (ASV) found in the Sediment enrichment. a) Prokaryotes. b) Eukaryotes.

## 4. Discussion

### 4.1. Sediment enrichment showed the highest activity and microbial synergy

Among the four enrichments, the Sediment enrichment exhibited the most promising methanotrophic and microalgal cooperation, maintaining high and stable activity throughout the incubation period. Marine sediments generally show higher methanotrophic activity than seawater or freshwater, due to factors such as a greater availability of CH_4_ from organic matter decomposition by methanogenic archaea, which supports a higher abundance of methanotrophs (Weber et al., 2019). Additionally, the aerobic conditions in sediments favor CH_4_ oxidation more than the static conditions in seawater and freshwater (He et al., 2019; Mao et al., 2022). Transitional coastal zones provide abundant nutrients and electron acceptors essential for methanotrophic activity. Sediment environments, in particular, offer a stable nutrient supply, with significantly higher levels of nitrate, ammonia, and phosphate compared to offshore waters, creating an ideal habitat for methanotrophs. This nutrient richness supports microbial life more effectively, leading to enhanced CH_4_ oxidation, increased photosynthetic microorganisms, and overall improved methanotrophic efficiency, as observed in the Sediment enrichment.

*Methylobacter marinus*/*whittenbury* emerged as the dominant methanotroph in the methalgae culture, likely due to selective enrichment under the provided conditions. This Type I methanotroph flourished in the MSM medium, which contained a high copper concentration (2.5 mg L^−1^ CuSO_4_·5H_2_O), essential for the function of particulate methane monooxygenase (pMMO). The copper-dependent pMMO enzyme, which oxidizes CH_4_ to methanol, is predominantly expressed in Type I methanotrophs under copper-rich conditions, while the soluble methane monooxygenase (sMMO) is expressed in copper-limited environments (Zhu et al., 2022). As a halotolerant species, *Methylobacter marinus*/*whittenbury* grows in salinities of 0.5 to 2 g NaCl L^−1^ (Bowman, 2006) and synthesizes ectoine as an osmoprotectant in high-salinity environments during CH_4_ bioconversion (Sahoo et al., 2021). Another species identified in the Sediment enrichment was *Methylophaga marina*, a halophilic obligate methylotroph that metabolizes one-carbon compounds like methanol. *Methylophaga marina* grows by utilizing by-products such as methanol, formaldehyde, or formate released by methanotrophs, as well as small carbohydrates and amino acids produced during photosynthesis (Dedysh and Dunfield, 2014; Ho et al., 2016).

The main photosynthetic microorganism identified in the sediment enrichment was *Picochlorum oklahomensis*, a broadly halotolerant chlorophyte microalga from the ‘pico’ size class (1–4 μm). Known for its small, oblong shape and cell wall, this microalga exhibits high biomass productivity and rapid growth, making it useful for biomass, pigment, and lipid production (Dahmen et al., 2014). *Picochlorum oklahomensis* is also valued for its high carotenoid content and resilience to a wide range of environmental conditions, including temperatures from 0 to 40 °C, pH variation, and salinities from 0 to 140 g L^−1^ (Zhu and Dunford, 2013).

In the Sediment enrichment, *Methylobacter marinus*/*whittenbury* and *Picochlorum oklahomensis* formed a synergistic community, likely driven by cross-feeding of metabolites. This interaction influenced both methylotrophic and non-methanotrophic bacteria, as no additional carbon sources were provided. *Methylophaga marina* likely utilized methanol or other C1 compounds produced by *Methylobacter marinus*/*whittenbury* during CH_4_ oxidation.

Additionally, non-methanotrophic heterotrophs, such as *Hyphomonas adhaerens* and *Labrenzia* sp., thrived by metabolizing organic byproducts like methanol, acetate, volatile fatty acids, and polysaccharides produced by methanotrophs and microalgae (Avila-Nuñez et al., 2024; Li et al., 2022; Zhang et al., 2023). Elevated heterotrophic richness in mixed cultures may stimulate methanotrophic activity through complex interactions, though the specifics remain unclear (Ho et al., 2014).

The Sediment enrichment, with the highest CH_4_ removal efficiency among the four enrichments, was deemed the most suitable for further experiments. Both *Methylobacter marinus*/*whittenbury* and *Methylophaga marina* are known ectoine producers. *Methylobacter marinus*/*whittenbury* can accumulate up to 5% ectoine of its dry cell weight in 4% NaCl (Eshinimaev et al., 2007), while *Methylophaga marina* can accumulate 15–19% in 5% NaCl (Doronina et al., 2010). The lower ectoine levels observed in this study’s Sediment enrichment may be due to dilution from the presence of microalgae and heterotrophic bacteria, which do not accumulate ectoine (Pérez et al., 2022).

### 4.2. High NaCl concentrations adversely affected methanotrophic activity

The NaCl concentrations above 3% significantly reduced CH_4_ RE, with experiments at 0% and 3% NaCl showing a 13-fold higher CH_4_ RE compared to those at 6% and 9% NaCl. This reduction might be linked to the limited halotolerance of the methanotrophic bacteria. Although there are reports indicating that these bacteria can tolerate saline concentrations up to 12% NaCl (Urakami and Komagata, 1987), such high salinity conditions are not ideal for growth, as the natural environment from which the samples were retrieved had a typical sea salt concentration of ∼3%. As a result, growth, abundance, and metabolic activity, including CH_4_ oxidation, were hindered at higher salinities (Ho et al., 2018). In contrast, the activity of the microalga *Picochlorum oklahomensis*, remained unaffected by varying salinities. As shown in **Fig. S6**, there was even net O_2_ production in some cases (e.g., comparing biological duplicates 9%A and 9%B), in contrast to the O_2_ consumption by bacteria. *Picochlorum oklahomensis* is highly adapted to a wide range of NaCl concentrations, up to 14%, and naturally thrives in hypersaline environments, allowing it withstand to significant fluctuations in salinity (Henley et al., 2004).

Despite the differences in the metabolic activities of methanotrophs and microalgae, the final biomass remained similar across all salinities, with only minor fluctuations in the relative abundances of each microorganism group. Under conditions of no salinity stress (0% NaCl) or at typical sea concentrations (3% NaCl), the CH_4_ removal efficiencies were consistent with those observed during the enrichment period. Microalgal growth was unaffected by increased salinity, maintaining similar abundances across all tests, except in the photoinhibited tests.

The methanotrophs’ response to salinity changes was most pronounced at 0% NaCl, where the decrease in their relative abundance could be attributed to their original marine environment, as natural methanotrophic populations from marine settings tend to be less resistant to reduced salinity (Osudar et al., 2017).

Ectoine was detected only in the initial inoculum and the 3% NaCl test, with concentrations ranging from 8.6 ± 0.03 to 12 ± 2.70 mg_ectoine_ g_VSS_^−1^. At higher salinities, ectoine was absent, likely because the ectoine present in the initial inoculum may have degraded. Ectoine synthesis requires significant carbon and nitrogen assimilation, which becomes unsustainable when metabolic activity is compromised by high salinity. This degradation could disrupt osmotic balance, but methanotrophs might recycle these nutrients as a last resort for energy (Reshetnikov et al., 2020). Cantera et al. (2016) also observed that a 6% NaCl concentration is not optimal for intracellular ectoine accumulation in methanotrophs, with even higher concentrations, such as 9%, resulting in lower ectoine yields.

To the authors’ knowledge, no chlorophyte microalgae have been confirmed to accumulate this osmolyte, which may have contributed to a dilution effect on the culture’s overall ectoine content. However, despite this potential dilution effect, the presence of microalgae reduced the need for external aeration by enhancing O_2_ production. This highlights the important role of microalgae in methanotrophic-microalgal cultures. Previous studies have shown that methanotrophs alone tend to rapidly become O_2_-limited, a problem that microalgae can help mitigate, especially when pure biogas will be considered as the main carbon source (Ruiz-Ruiz et al., 2020; Van Der Ha et al., 2011).

### 4.3. Moderate salinity shock boosted ectoine levels without compromising methanotrophic activity

Similar to the salinity test, a 6% NaCl concentration significantly reduced the CH_4_ RE, lowering it to levels 4 to 6 times below the initial values, depending on whether the starting salinity was 4.5% or 6% NaCl (**Fig. 4a**). This reduction can be linked to the limited halotolerance of methylotrophic bacteria (Urakami and Komagata, 1987). Additionally, the short intervals between osmotic shocks may have further impacted methanotroph activity. Frequent osmotic shocks lasting 24 h or less can induce stress that either enhances or inhibits CH_4_ oxidation, depending on the specific methanotroph species present (Osudar et al., 2017).

Regarding biomass concentration, there was no clear pattern in the relationship between osmotic shocks and biomass levels. This may be due to significant changes in the microbial community composition within the mixed culture caused by the osmotic shocks. Some bacterial species may become more dominant, while less adaptable species may decline following sudden changes in salinity (Bissett et al., 2012). Given the rapid growth rates of these microorganisms, such shifts can occur abruptly within just 24 h after each osmotic shock (Kwon et al., 2019; Zhu and Dunford, 2013). Methanotrophs typically respond less severely to gradual increases in salinity than to sudden osmotic shocks, as gradual changes allow more time for physiological and gene expression adjustments (Han et al., 2017; Osudar et al., 2017). This gradual adaptation may explain why sudden osmotic shocks, particularly at higher salinities, have a more pronounced negative effect on CH_4_ oxidation and overall microbial activity.

In the three tests, more than 99% of the biomass consisted of bacteria (methanotrophic and heterotrophic), with almost no detectable microalgae. This can be explained to the small relative abundance of microalgae in the initial inoculum, which contained only 0.03% microalgae (**Table S1**). The decline in microalgae abundance compared to the salinity tests can be attributed to the slower growth rates of microalgae relative to methanotrophs and heterotrophic bacteria. Methanotrophic bacteria have growth rates greater than 0.1 h^−1^, with a doubling time of 18 h (Amaral and Knowles, 1995; Kwon et al., 2019), while *Picochlorum oklahomensis* has a slower growth rate of 0.5 to 0.9 day^−1^ and a doubling time of 1.4 days, depending on salinity (Foflonker et al., 2018; Zhu and Dunford, 2013). This difference in growth rates and doubling times, along with the initial biomass ratio, which affects carbon uptake, CH_4_ oxidation efficiencies, and biomass growth (Ruiz-Ruiz et al., 2020), contributed to the dominance of bacteria in the culture.

The highest ectoine concentration, reaching 51.3 ± 1.1 mg_ectoine_ g_VSS_^−1^, was obtained when the NaCl content was increased from 3% to 4.5%. This result is consistent with reports of ectoine production by other methanotrophic bacteria (**Table 2**) and surpasses the concentrations observed in the salinity test. The higher ectoine levels are likely due to the greater abundance of bacteria in the culture (nearly 100%) compared to the 30–50% observed during salinity tests. Although microalgae in the culture can dilute ectoine concentration, they play a crucial role in enhancing the system’s sustainability (Ruiz-Ruiz et al., 2020).

**Table 2.** Specific ectoine values in reported methanotrophic cultures and this study.

| Methanotrophic organism | Specific ectoine content | Reference |
| --- | --- | --- |
| *Methylomicrobium alcaliphilum* 20Z | $66.9 \pm 4.2 \text{ mg}_{\text{ectoine}} \text{ g}_{\text{TSS}}^{-1}$ | (Cantera et al., 2016) |
| *Methylomicrobium alcaliphilum* 20Z | $37.4 \pm 3.8 \text{ mg}_{\text{ectoine}} \text{ g}_{\text{TSS}}^{-1}$ | (Cantera et al., 2017b) |
| *Methylomicrobium alcaliphilum* 20Z | $94.2 \pm 10.1 \text{ mg}_{\text{ectoine}} \text{ g}_{\text{TSS}}^{-1}$ | (Cantera et al., 2018) |
| Enrichment of haloalkaliphilic bacteria | $79.7 \pm 5.1 \text{ mg}_{\text{ectoine}} \text{ g}_{\text{TSS}}^{-1}$ | |
| *Methylomicrobium alcaliphilum* 20Z | $20.7 \pm 1.2 \text{ mg}_{\text{ectoine}} \text{ g}_{\text{wet biomass}}^{-1}$ | (Mustakhimov et al., 2019) |
| *Methylomicrobium alcaliphilum* 20Z | $109 \text{ mg}_{\text{ectoine}} \text{ g}_{\text{TSS}}^{-1}$ | (Cantera et al., 2020) |
| Enriched haloalkaliphilic consortium | $3.0 \pm 0.6 \text{ mg}_{\text{ectoine}} \text{ m}^{-3} \text{ h}^{-1}$ | |
| Enriched haloalkaliphilic consortium | $56.6 \pm 2.5 \text{ mg}_{\text{ectoine}} \text{ g}_{\text{VSS}}^{-1}$ | (Carmona-Martínez et al., 2021) |
| *Methylomicrobium alcaliphilum* 20Z | $111 \text{ mg}_{\text{ectoine}} \text{ g}_{\text{TSS}}^{-1}$ | (Cho et al., 2022) |
| Consortium from salt lagoon | $30 \pm 4 \text{ mg}_{\text{ectoine}} \text{ g}_{\text{VSS}}^{-1}$ | (Rodero et al., 2022) |
| Consortium from salt lagoon | 20-52 mg<sub>ectoine</sub> g<sub>VSS</sub><sup>-1</sup> | (Rodero et al., 2023) |
| Mixed methanotrophic culture | 79 mg<sub>ectoine</sub> g<sub>VSS</sub><sup>-1</sup> | (Rodero et al., 2024) |
| Enrichment from marine coast sediment<br>(*Methylobacter marinus/whittenbury*) | 51.3 ± 1.1 mg<sub>ectoine</sub> g<sub>VSS</sub><sup>-1</sup> | **This study** |

The high abundance of heterotrophic and methanotrophic bacteria played a significant role in the overall ectoine content of the biomass in response to osmotic shocks. Ectoine levels increased with salinities up to 4.5% NaCl, but decreased under more extreme saline conditions (6% NaCl). This decline at higher salinities is likely due to physiological stress and potential cellular damage caused by severe osmotic pressure, which can disrupt cellular functions. Under such stress, bacteria may shift from accumulating ectoine to breaking it down for use as a carbon and nitrogen source. This dual function of ectoine, serving both as an osmoprotectant and a potential energy source, explains the observed decrease in ectoine levels under extreme osmotic conditions (Cantera et al., 2017b; Carmona-Martínez et al., 2021).

Unexpectedly low initial intracellular ectoine concentrations were observed during the early stages of experiments at 3% salinity and following the first osmotic shock. Typically, ectoine levels in the biomass remained above 6.7 ± 1.4 mg_ectoine_ g_VSS_^1^; however, in these instances, the values ranged from 1.91 ± 0.1 to 4.4 ± 0.2 mg_ectoine_ g_VSS_^−1^. These unusually low initial concentrations may be due to the centrifugation process used during mass sedimentation for medium replacement, which took approximately 2.5 hours. According to Cantera et al. (2017a), methanotrophs can be adversely affected by mechanical agitation, which may impair their ability to produce or maintain ectoine levels. When methanotrophs are subjected to mechanical stress, key enzymes encoded by the gene cluster *doeBDAC* are activated, enabling the degradation of ectoine and its integration into their metabolism (Reshetnikov et al., 2020). This process helps conserve nitrogen and carbon, while also supporting energy conservation during stressful conditions. This mechanism is evident in the significant decrease in ectoine levels following the increase in salinity during the second osmotic shock in OS1, as well as the lack of increase in biomass concentration under sustained stress in OS2 and OS3 (**Fig. 4b**). This adaptive response allows methanotrophs to dynamically adjust to environmental changes, particularly in saline conditions, which is crucial for their survival and efficiency in utilizing CH_4_ as a carbon source (Reshetnikov et al., 2020).

## 5. Conclusions

This study demonstrates the potential of methanotrophic-microalgal cultures to valorize biogas by converting CH_4_ and CO_2_ into the high-value compound ectoine. Moderate NaCl concentrations enhanced ectoine production, whereas higher salt levels hindered both CH_4_ removal and ectoine accumulation. Osmotic shocks revealed the culture’s resilience, with methanotrophic bacteria adapting to moderate salinity changes, but struggling at higher concentrations. The cultivation of methanotrophic bacteria with microalgae provided an ecological advantage by utilizing CO_2_, often overlooked in methanotroph-only systems, with the potential of boosting O_2_ production to sustain CH_4_ oxidation. This is the first study to explore ectoine production via the methalgae approach.

## CRediT authorship contribution statement

**Patricia Ruiz-Ruiz**: Conceptualization, Data curation, Formal analysis, Investigation, Methodology, Writing original draft, review and editing. **Patricia Mohedano-Caballero**: Formal analysis, Investigation, Data curation. **Jo De Vrieze**: Conceptualization, Funding acquisition, Project administration, Resources, Supervision, Writing – review and editing.

## Funding

This research was funded by the Research Foundation of Flanders (FWO Project Bio-Economy "EctoMet" G0E0723N).

## Acknowledgements

The authors express their gratitude to the Agency for Nature and Forest in the province of West Flanders for providing sampling authorization. Special thanks are extended to forest ranger Koen Maréchal for his invaluable support during the sampling process. The authors also wish to acknowledge Jana De Bodt, Greet Van de Velde, and Tim Lacoere for their technical assistance and support with the cultures, equipment, and molecular analysis.

## Notes

### Competing Interest Statement

The authors have declared no competing interest.

