## Supplementary Information for "Ectoine production through a marine methanotroph-microalgae culture allows complete biogas valorization"

### Supplementary Figures

Zwin Natuurpark
Knokke-Heist

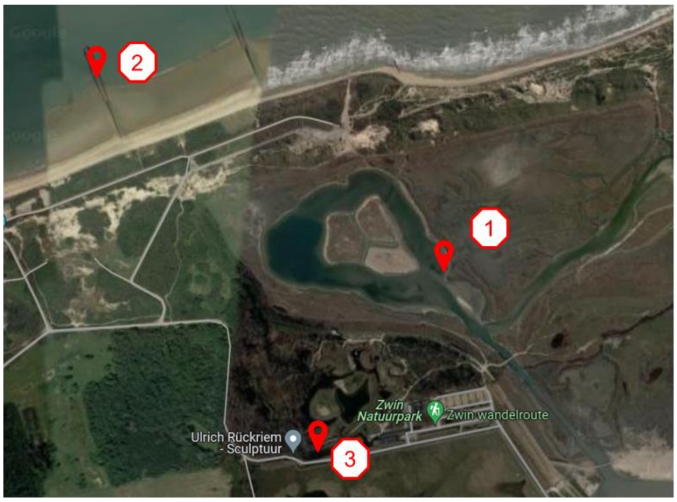


(1) 51°21'38.0"N 3°21'09.8"E pH 7,43

(2) 51°21'56.6"N 3°20'21.9"E pH 7,86

(3) 51°21'28.9"N 3°20'45.9"E pH 7,70

**Fig. S1.** Sampling points at the Zwin Natuur Park (Google maps).

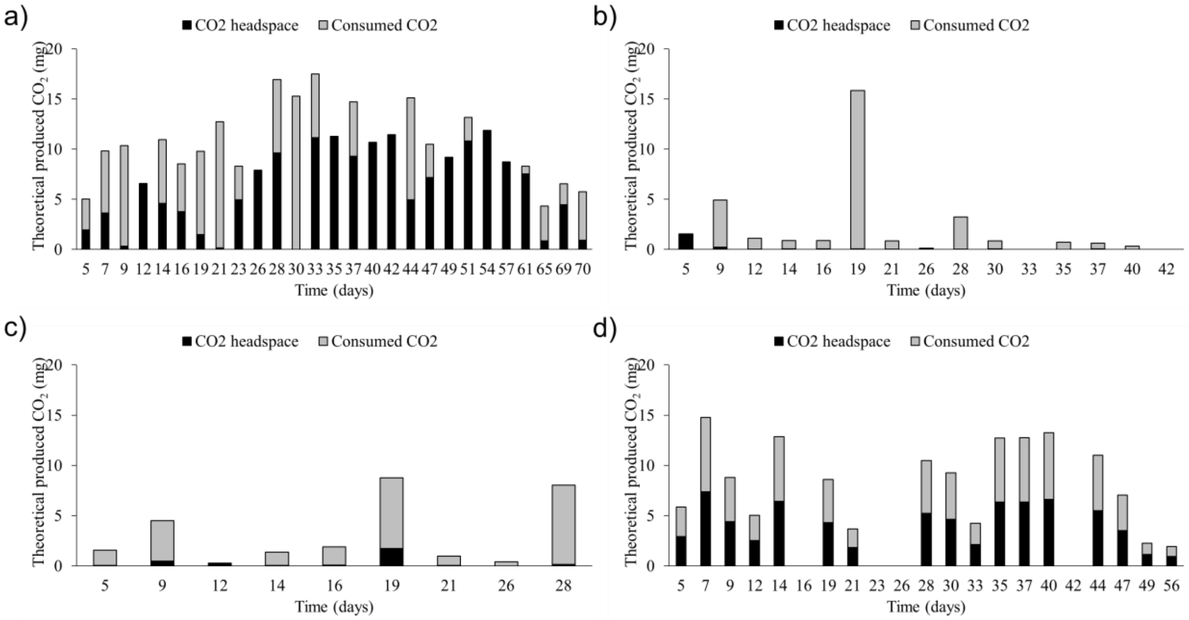


**Fig. S2.** Theoretically produced $CO_2$ from $CH_4$ consumption, from which: consumed $CO_2$ (black), and remaining $CO_2$ in the headspace (grey)*. a) Sediment. b) River. c) North Sea. d) Combination.

***Theoretical calculations of $CO_2$ production by methanotrophs:**

Stoichiometric reaction by methanotrophs:

$CH_4 + 2O_2 \rightarrow CO_2 + 2H_2O$

This reaction indicates that 1 mol of methane ($CH_4$) produces 1 mol of carbon dioxide ($CO_2$).

Mass calculation for $CO_2$ produced from $CH_4$ consumption:

- The molar mass of methane ($CH_4$) is 16.04 g/mol.
- The molar mass of carbon dioxide ($CO_2$) is 44.01 g/mol.
- The mass conversion ratio is calculated as follows:

$$Mass\ convertion\ ratio = \frac{Molar\ mass\ of\ CH_4}{Molar\ mass\ of\ CO_2} = \frac{16.04\ g/mol}{44.01\ g/mol} \approx 2.744$$

According to Leak and Dalton (1986), Type I methanotrophs exhibit a yield ratio of 0.6 for $CO_2$ production per $CH_4$ consumed. The theoretical mass of $CO_2$ produced can therefore be calculated using the formula:

$$Theoretical\ CO_2\ (in\ mg) = (Consumed\ mass\ of\ CH_4\ (in\ mg) \times 2.7444) \times 0.6$$

**Net $CO_2$ consumption calculation:**

To determine the net amount of $CO_2$ consumed, the mass of $CO_2$ remaining in the headspace was substracted from the theoretical mass of $CO_2$ produced:

$$Consumed\ CO_2\ (in\ mg) = Theoretical\ CO_2\ (in\ mg) - \ CO_2\ remaining\ in\ headspace\ (in\ mg)$$

For mass calculations, headspace volume, a temperature of 28 °C, and an atmospheric pressure of 1 atm were considered.

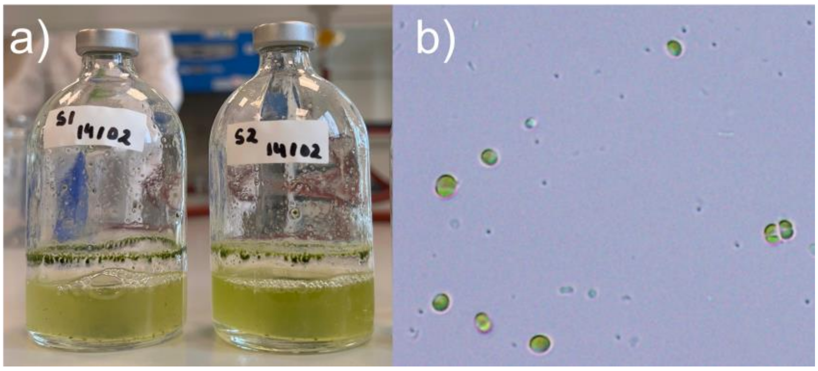


**Fig. S3.** a) Sediment enrichment bottles with visible microalgal growth. b) Light microscopy picture of the Sediment enrichment (20X).

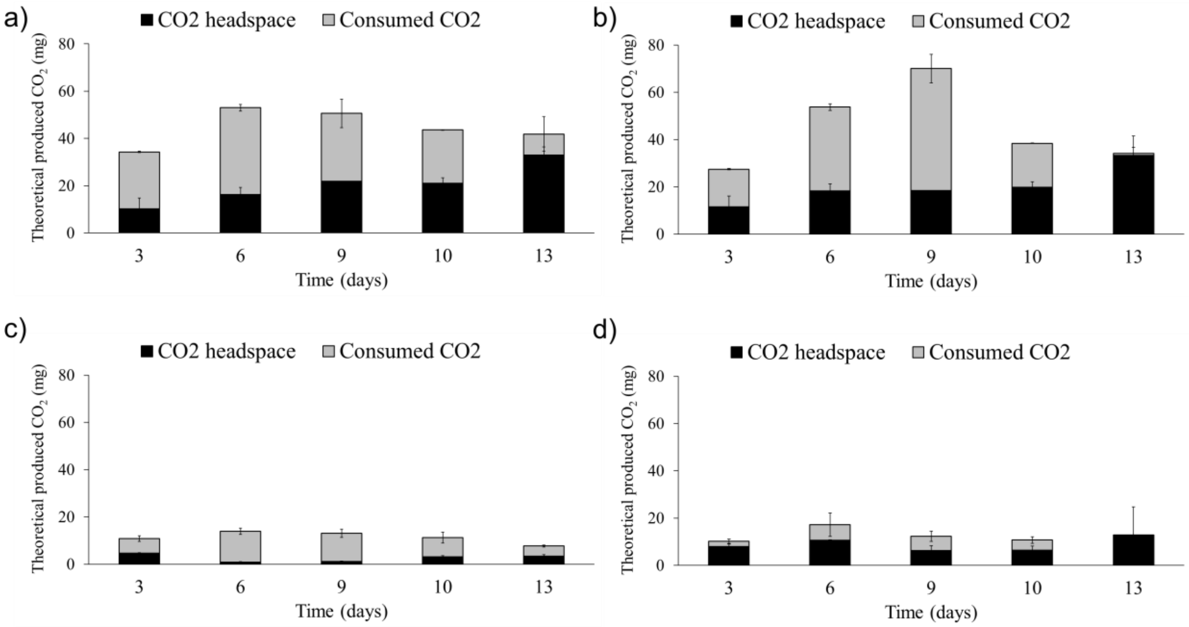


**Fig. S4.** Theoretically produced $CO_2$ from $CH_4$ consumption, from which: consumed $CO_2$ (black), and remaining $CO_2$ in the headspace (grey). a) 0% NaCl test. b) 3% NaCl test. c) 6% NaCl test. d) 9% NaCl test. Error bars indicate the standard deviation of biological duplicates.

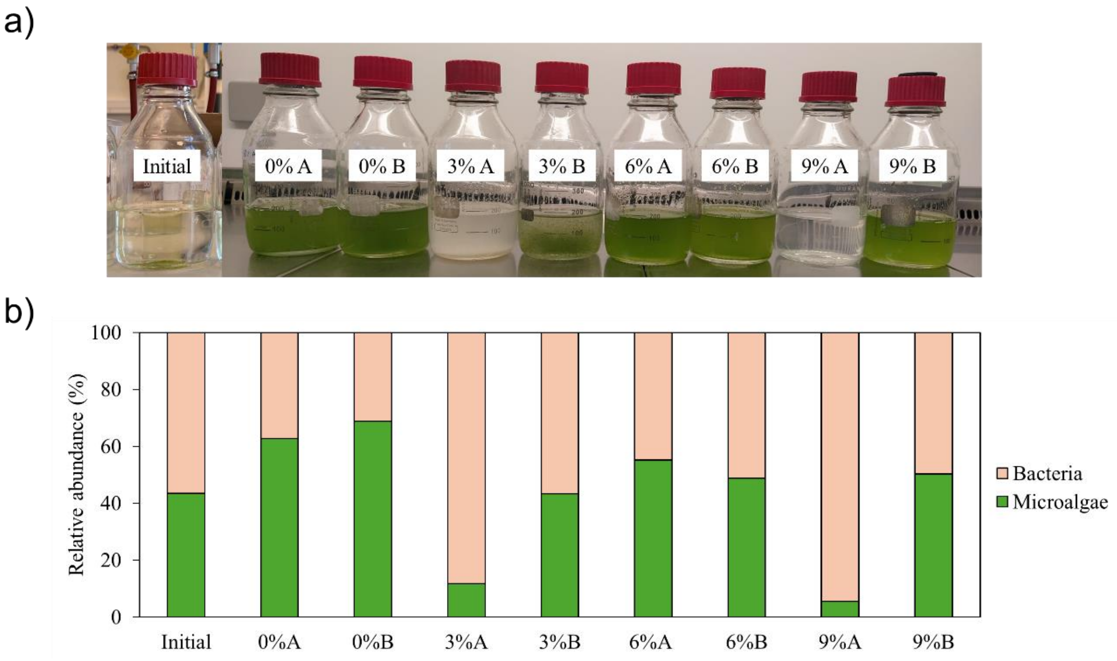


**Fig. S5.** a) Photoinhibition on 3A and 9A reactors. b) Relative abundance of initial and final bacterial and microalgal cells measured by FCM during the NaCl concentration tests.

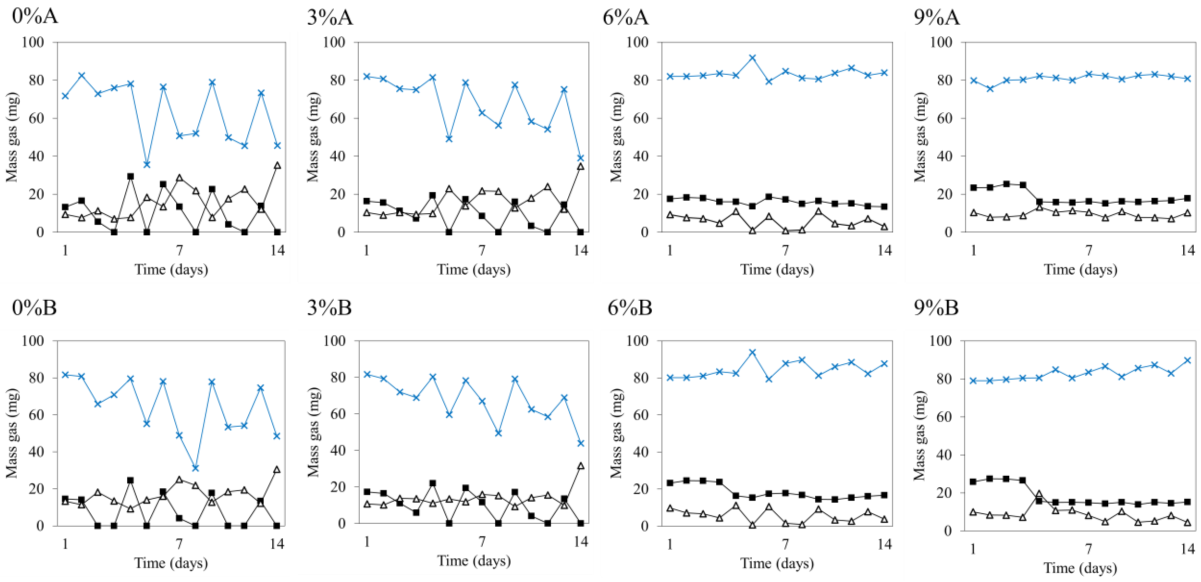


**Fig. S6.** Gas evolution in the headspace for each biological duplicate at different NaCl concentrations. Black squares indicate $CH_4$, white triangles $CO_2$, and blue crosses $O_2$.

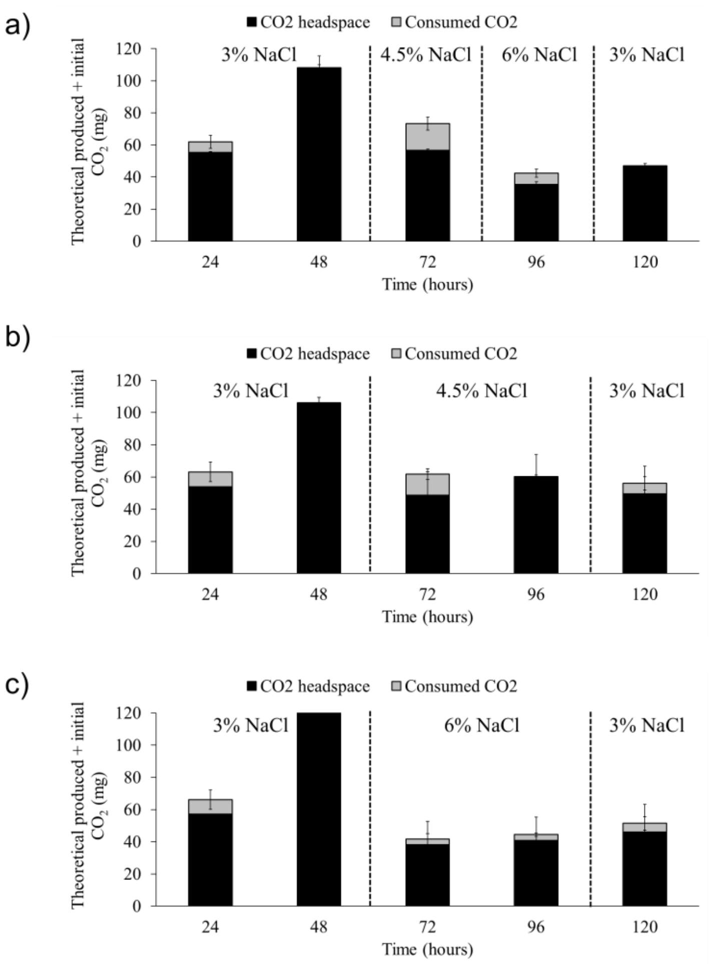


**Fig. S7.** Theoretically produced $CO_2$ from $CH_4$ consumption + initial $CO_2$, from which: consumed $CO_2$ (black), and remaining $CO_2$ in the headspace (grey). a) OS1 test. b) OS2 test. c) OS3 test. Error bars indicate the standard deviation of biological duplicates.

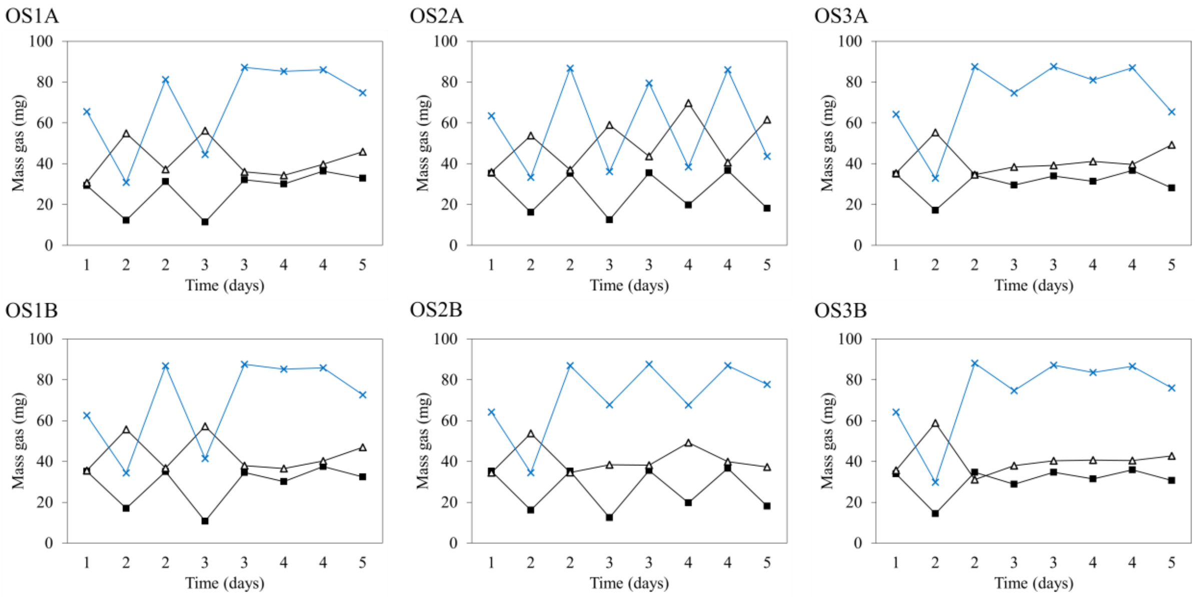


**Fig. S8.** Gas evolution in the headspace for each biological duplicate for OS1, OS2 and OS3. Black squares indicate $CH_4$, white triangles $CO_2$, and blue crosses $O_2$.

**Table S1.** Percentage of bacteria and microalgae obtained by FCM during the osmotic shocks for OS1, OS2, OS3.

| | OS1 | | OS2 | | OS3 | |
|---|---|---|---|---|---|---|
| | Microalgae (%) | Bacteria (%) | Microalgae (%) | Bacteria (%) | Microalgae (%) | Bacteria (%) |
| Initial 3% NaCl | 0.03 | 99.97 | 0.03 | 99.97 | 0.03 | 99.97 |
| Final 3% NaCl | 0.00 | 100.00 | 0.00 | 100.00 | 0.00 | 100.00 |
| Initial 1st shock | 0.03 | 99.97 | 0.01 | 99.99 | 0.04 | 99.96 |
| Final 1st shock | 0.00 | 100.00 | 0.00 | 100.00 | 0.00 | 100.00 |
| Initial 2nd shock | 0.09 | 99.91 | 0.00 | 100.00 | 0.00 | 100.00 |
| Final 2nd shock | 0.00 | 100.00 | 0.00 | 100.00 | 0.00 | 100.00 |
| Initial after shock | 0.06 | 99.94 | 0.03 | 99.97 | 0.06 | 99.94 |
| Final after shock | 0.00 | 100.00 | 0.00 | 100.00 | 0.00 | 100.00 |
